# Optogenetically induced low-frequency correlations impair perception

**DOI:** 10.1101/252841

**Authors:** Anirvan S. Nandy, Jonathan J. Nassi, John H. Reynolds

## Abstract

Deployment of covert attention to a spatial location can cause large decreases in low-frequency correlated variability among neurons in macaque area V4 whose receptive-fields lie at the attended location. It has been estimated that this reduction accounts for a substantial fraction of the attention-mediated improvement in sensory processing. These estimates depend on assumptions about how population signals are decoded and the conclusion that correlated variability impairs perception, is purely hypothetical. Here we test this proposal directly by optogenetically inducing low-frequency fluctuations, to see if this interferes with performance in an attention-demanding task. We find that low‐frequency optical stimulation of neurons in V4 elevates correlations among pairs of neurons and impairs the animal’s ability to make fine sensory discriminations. Stimulation at higher frequencies does not impair performance, despite comparable modulation of neuronal responses. These results support the hypothesis that attention-dependent reductions in correlated variability contribute to improved perception of attended stimuli.

## INTRODUCTION

Neurons exhibit responses that are highly variable (Shadlen and Newsome, 1998), with nearby neurons in the cortex exhibiting correlated variability in their spiking output (Cohen and Kohn, 2011; Smith and Kohn, 2008; Smith and Sommer, 2013; Zohary et al., 1994). It has been estimated on theoretical grounds that even weak correlations substantially reduce the information coding capacity of a population (Zohary et al., 1994). Spatial attention can reduce correlated variability (often referred to as noise correlations) among neurons in macaque visual area V4 (Cohen and Maunsell, 2009; Mitchell et al., 2009), an area that is strongly modulated by spatial attention (Reynolds and Chelazzi, 2004). Mitchell et al. (2009) found that this reduction is restricted to low frequencies below 10Hz. These studies have estimated, on theoretical grounds, that the reduction in correlated variability accounts for a large fraction (about 80%) of the perceptual benefit due to attention. However, these estimates rely on specific assumptions about the relationship between noise and signal correlations, and thereby, on how population signals are read out in the brain (Averbeck et al., 2006; Moreno-Bote et al., 2014). Theoretical studies using heterogeneous tuning curves and optimal readout have concluded that correlated variability does not necessarily limit information (Ecker et al., 2011; Shamir and Sompolinsky, 2006). Consistent with the interpretation that attention-dependent reductions in correlated variability improve perception, a recent study of the effects of naturally occurring fluctuations in neural correlations found improved sensory discrimination when neurons in Area V4 were desynchronized (Beaman et al., 2017). However, other studies have posited that correlations themselves may be induced by fluctuations in attention (Goris et al., 2014), resulting in variation in response gain that is shared across neurons, and other experiments have shown that attention can, under some conditions, also increase correlated neural response variability (Ruff and Cohen, 2014). Taken together, these studies call into question the simple idea that attention reduces correlation so as to improve sensory discrimination. Importantly, all of these studies are correlative in nature. The causal role of correlated variability in perception has not been tested and thus the proposal that low-frequency correlated variability is detrimental for perception has remained purely hypothetical.

Here, we sought to directly test the effects of correlated variability on sensory discrimination by using optogenetic activation to induce correlations in Area V4 as monkeys performed an orientation discrimination task near perceptual threshold. We exploited the fact that attentional modulation of correlated variability is both spatially‐ and frequency selective: attention-dependent reductions in correlation are restricted to low frequencies (<10 Hz (Mitchell et al., 2009)). We reasoned that the correlations that impair perception may have an inherent time scale, with low‐ but not high-frequency correlations impairing perception. If so, we would predict that the effects of correlations on perception should be specific to this low-frequency range.

## RESULTS AND DISCUSSION

We took advantage of a novel approach to primate optogenetics and electrophysiology (Nassi et al., 2015; Ruiz et al., 2013) in which the native dura mater is replaced by a silicone based artificial dura (Fig 1A,B). This approach provides an optically clear window into the awake-behaving primate brain and allows precise opto‐electrophysiology. We injected a lenti-viral construct (lenti-CaMKIIa-C1V1E162T-ts-EYFP) to preferentially drive expression of the depolarizing opsin C1V1 in excitatory neurons (Han et al., 2009) in a restricted portion (200-300*μm* diameter) of dorsal V4 of two macaque monkeys (1C). Despite some heterogeneity in orientation tuning width at each injection site, overall there was similar tuning among neurons within a site (Fig 2 - Supp 1).

**Fig 1.**
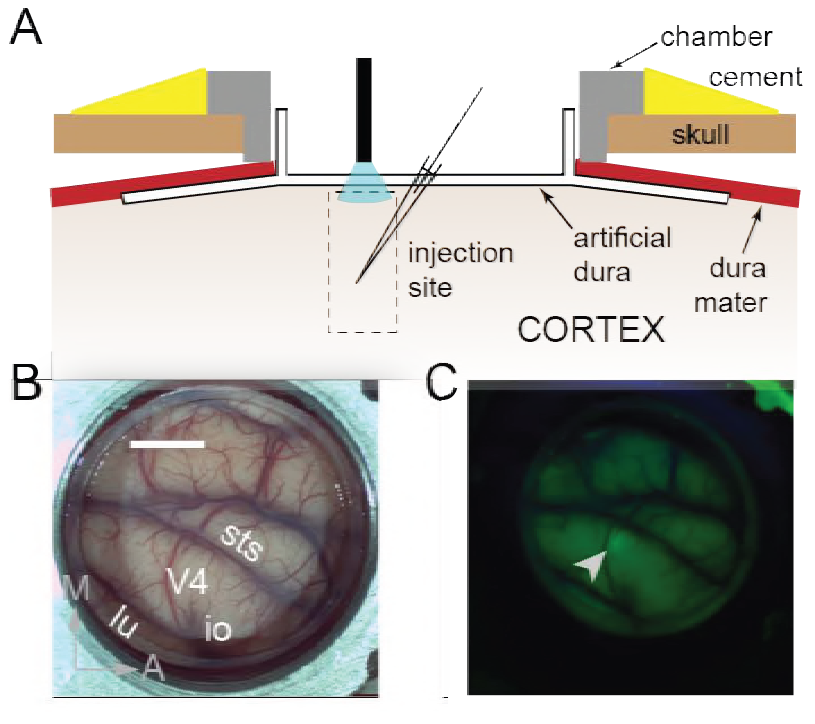
Surface Optogenetics and electrophysiology through an artificial dura. (**A**) Schematic of an artificial dura (AD) chamber. A portion of the native dura mater (red) is resected and replaced with a silicone based optically clear artificial dura (AD). The optical clarity of the AD allows precisely targeted injections of viral constructs and subsequent optical stimulation and electrophysiological recordings. (**B**) An AD chamber is shown over dorsal V4 in the right hemisphere of Monkey A. sts = superior temporal sulcus, lu = lunate sulcus, io = inferior occipital sulcus. Area V4 lies on the pre-lunate gyrus between the superior temporal and lunate sulci. Scale bar = 5mm; M=medial, A=anterior (**C**) EYFP expression at the first injection site (lenti-CaMKIIα-C1V1-ts-EYFP) after 4 weeks.

We trained two monkeys to perform an attention-demanding orientation-change detection task (2A). The monkeys were spatially cued to attend to one of two spatial locations. In the ‘attend in’ condition, the monkeys were instructed to covertly attend to a spatial location within the receptive fields of neurons at the viral injection site, while maintaining fixation at a central fixation point. In the ‘attend-away’ condition attention was directed to a location of equal eccentricity across the vertical meridian. On each trial, a sequence of oriented Gabor stimuli simultaneously flashed on and off at both spatial locations (200ms on, variable 200-400ms off). At an unpredictable time (minimum 1s, maximum 5s), one of the two stimuli (95% probability at cued location; 5% probability at uncued location, ‘foil trials’) briefly changed in orientation (200ms) and the monkey was rewarded for making a saccade to the location of orientation change. If no change occurred within 5s (‘catch trials’, 13% of trials), the monkey was rewarded for holding fixation. We controlled task difficulty by varying the degree of orientation change and thereby obtained behavioral performance curves (psychometric functions) for each recording session (Fig 2B, Fig 2 - Supp 2A,B). Impaired performance (Fig 2 - Supp 2A, left panel, square symbol) and slower reaction times (Fig 2 - Supp 2A, right panel, square symbol) were observed for the foil trials, indicating that the monkey was indeed using the spatial cue in performing the task

**Fig 2.**
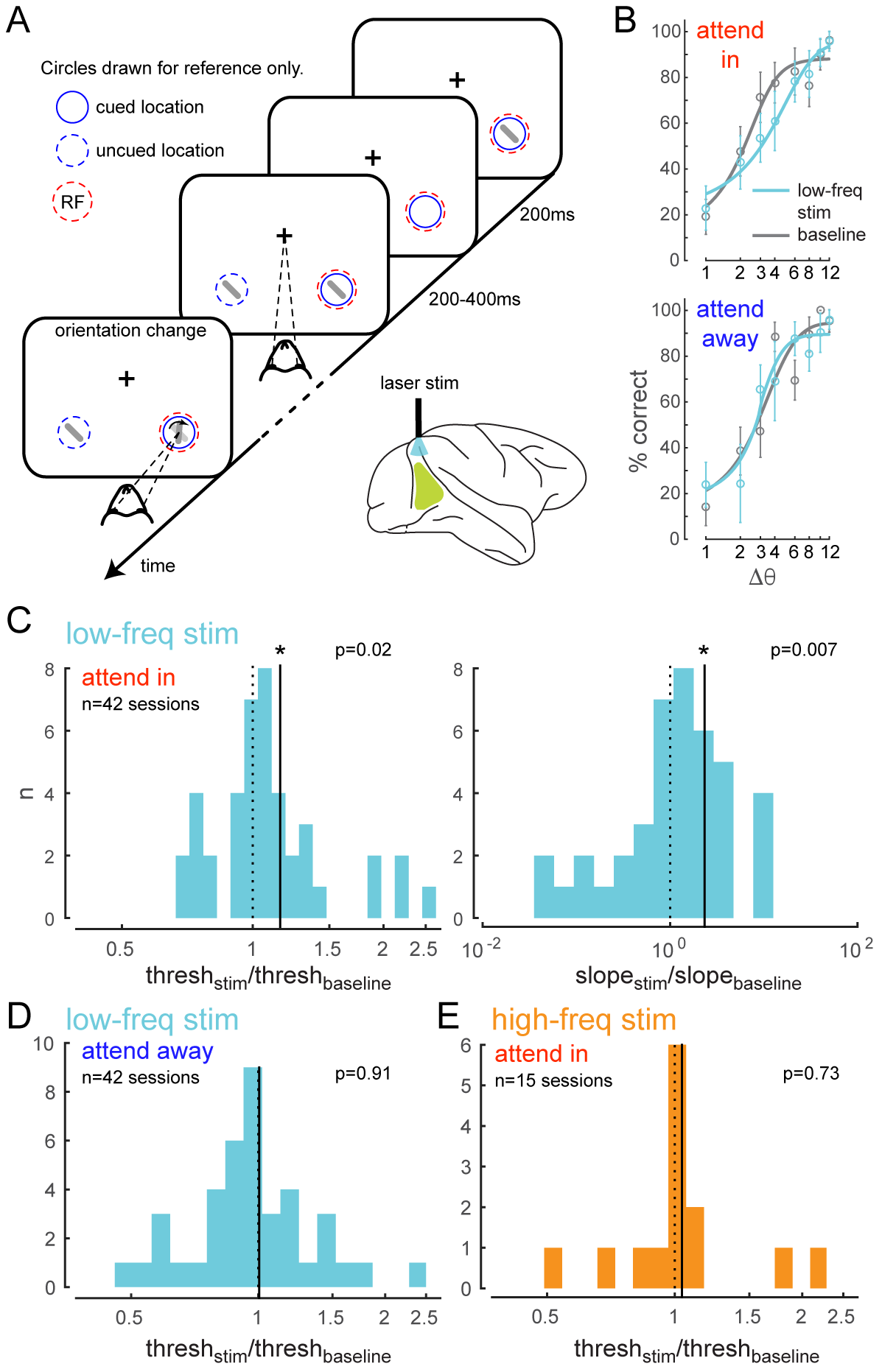
Optogenetically-induced low-frequency correlations cause a frequency‐ and spatially‐ selective impairment in an attention-demanding orientation discrimination task. (**A**) Attention task: While the monkey maintained fixation, two oriented Gabor stimuli (schematized as oriented bars) flashed on and off simultaneously at two spatial locations: one at the RF of the opsin injection site, the other at a location of equal eccentricity across the vertical meridian. The monkey was cued to covertly attend to one of the two locations. At an unpredictable time, one of the two stimuli changed in orientation. The monkey was rewarded for making a saccade to the location of orientation change at either location (95% probability of change at cued location; 5% probability at un-cued location [foil trials]). If no change occurred (catch trials), the monkey was rewarded for maintaining fixation. On a random subset of trials, the opsin site was optically stimulated using a low-frequency (4-5 Hz) sinusoidally modulated laser light (λ = 532nm). (**B**) Psychometric functions for an example behavioral session showing performance (hit rate) as a function of task difficulty (size of orientation change) for the baseline (no optical stimulation) condition in gray and low-frequency optical stimulation condition in blue. *Top*, monkey was instructed to attend to the site of optical stimulation; *Bottom*, monkey was instructed to attend to the contralateral hemifield. Error bars are std. dev. obtained by a jackknife procedure and corrected for the number of jackknives (20). The data has been fitted with a smooth logistic function. (**C**) Change in psychometric function threshold (left panel) and slope (right panel) due to low-frequency optical stimulation when the monkey was attending in to the site of optical stimulation across all behavioral sessions. Changes are expressed as a ratio over baseline. The solid line represents the mean of the distribution. Both changes are statistically significant. (**D-E**) No significant change in threshold either when the monkey was attending away from the site of optical stimulation (**D**) or due to high-frequency optical stimulation (**E**).

To test if low-frequency correlations impair discrimination we optically stimulated neurons at the opsin injection site with 4-5Hz sinusoidally modulated low-power laser stimulation, on a randomly chosen subset of trials (‘low-frequency stimulation’ condition). Our goal was to induce correlations without significantly altering the mean firing rates by using low-power stimulation. Significant changes in mean firing rate could have unknown effects such as masking of the stimulus evoked response. Equating firing rates also avoids any indirect effects of mean firing rate changes on spike-count correlations (Cohen and Kohn, 2011). We find that low-frequency optical stimulation modulates the timing of the neural response (4D) but does not alter the overall magnitude of the population response (Fig 4A,B,C). We replicate previous findings that attention reduces low-frequency spike-count correlations in the baseline (no optical stimulation) condition (Mitchell et al., 2009) (Fig 3A, left panel; gray versus white bars, *p* = 0.02, *t*-test). As predicted, low-frequency optical stimulation increases low‐ frequency correlations (Fig 3A, left panel; blue versus gray bar, *p* = 0.045, *t*-test). The induced correlations were at a level comparable in strength to that observed when attention was directed away from the RF location in the baseline condition (Fig 3A, left panel; blue versus white bar). Optogenetic activation is accompanied by a period of reduced activity following stimulation. By careful titration of laser intensity (amplitude of sinusoidal modulation) we were able to alter the timing of spiking without altering mean firing rate. This is shown in Figure 4: we see a robust increase in firing rate due to attention in both the low-frequency stimulation and baseline conditions (Fig 4A,B), but there is no significant rate increase due to optical stimulation either during the prestimulus blank period (Fig 4C, left panel, *p* = 0.49, *t*-test) or during the stimulus presentation period (Fig 4C, right panel, *p* > 0.1, *t*-test). Rather, unit activity shows phase locking to optical stimulation (Fig 4D,E). The distributions of spiking activity with respect to the phase of the optical stimulation show significant deviation from what would be expected from a null distribution (Fig 4D, example units, *p <<* 0.01; Fig 4E, population, *p* < 0.01, Rayleigh test). The null distributions were derived from a rate-Peri-matched Poisson process. Thus, the physiology data shows that we successfully induced correlated activity among neurons at the opsin site without affecting the response rates.

**Fig 3.**
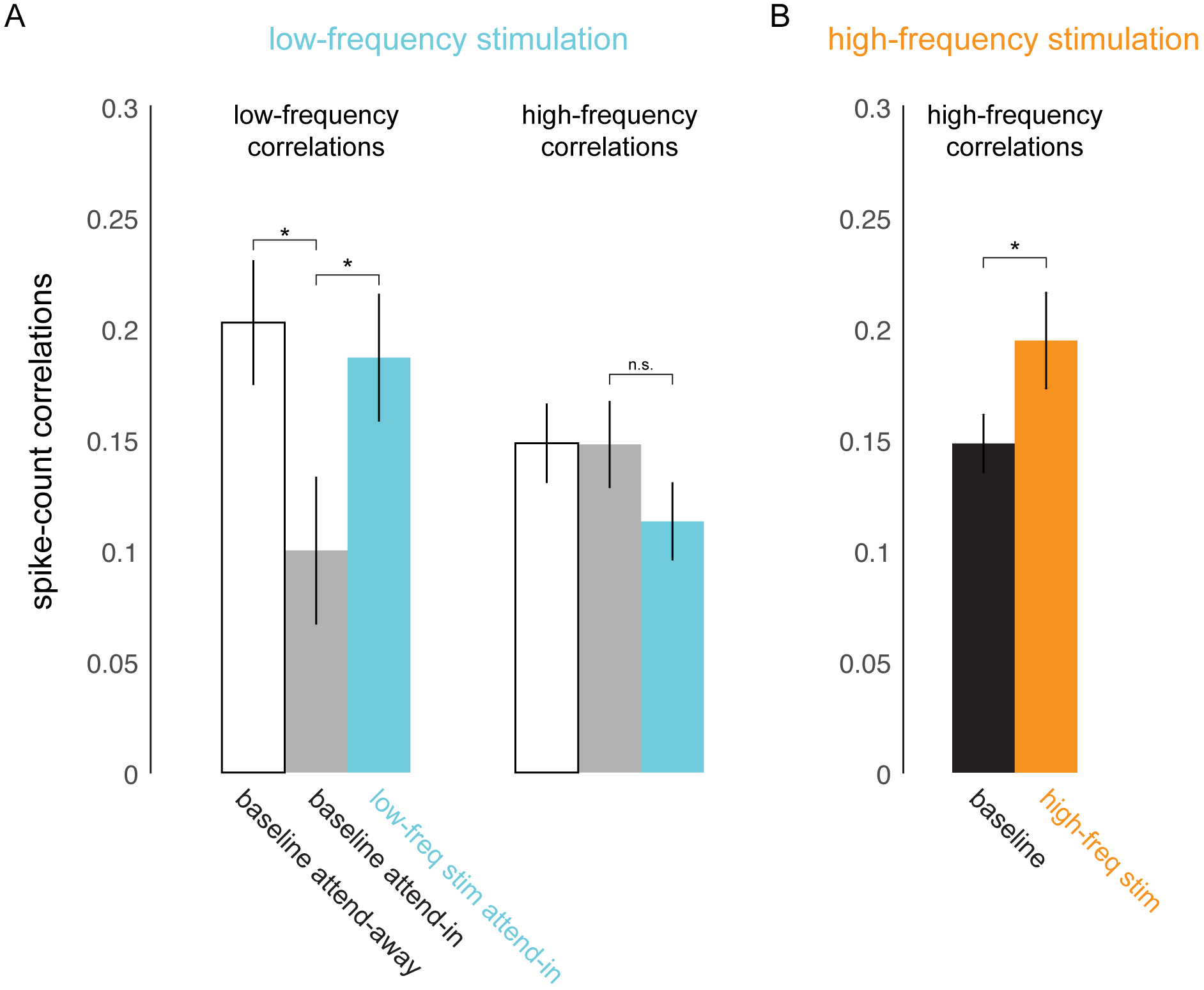
Optical stimulation at low‐ and high-frequencies induces lowy‐ and high-frequency correlated activity. (**A**) Consistent with earlier reports (Mitchell et al., 2009), attention reduces baseline spike-count correlations at low frequencies (200ms counting window, *p* = 0.02; left panel, white versus gray bar) but not at high frequencies (50ms window; right panel, white versus gray bar). Low-frequency optical stimulation increases low-frequency correlations (*p* = 0.045; left panel, gray versus blue bar) but not high‐frequency correlations (*p* > 0.1; right panel, gray versus blue bar). (**B**) High-frequency optical stimulation increases high-frequency correlations (*p* < 0.05). *n* = 79 pairs for baseline and low-frequency stimulation, *n* = 27 pairs for high-frequency stimulation, collapsed across attention conditions. Mean +/− s.e.m. in all plots.

**Fig 4.**
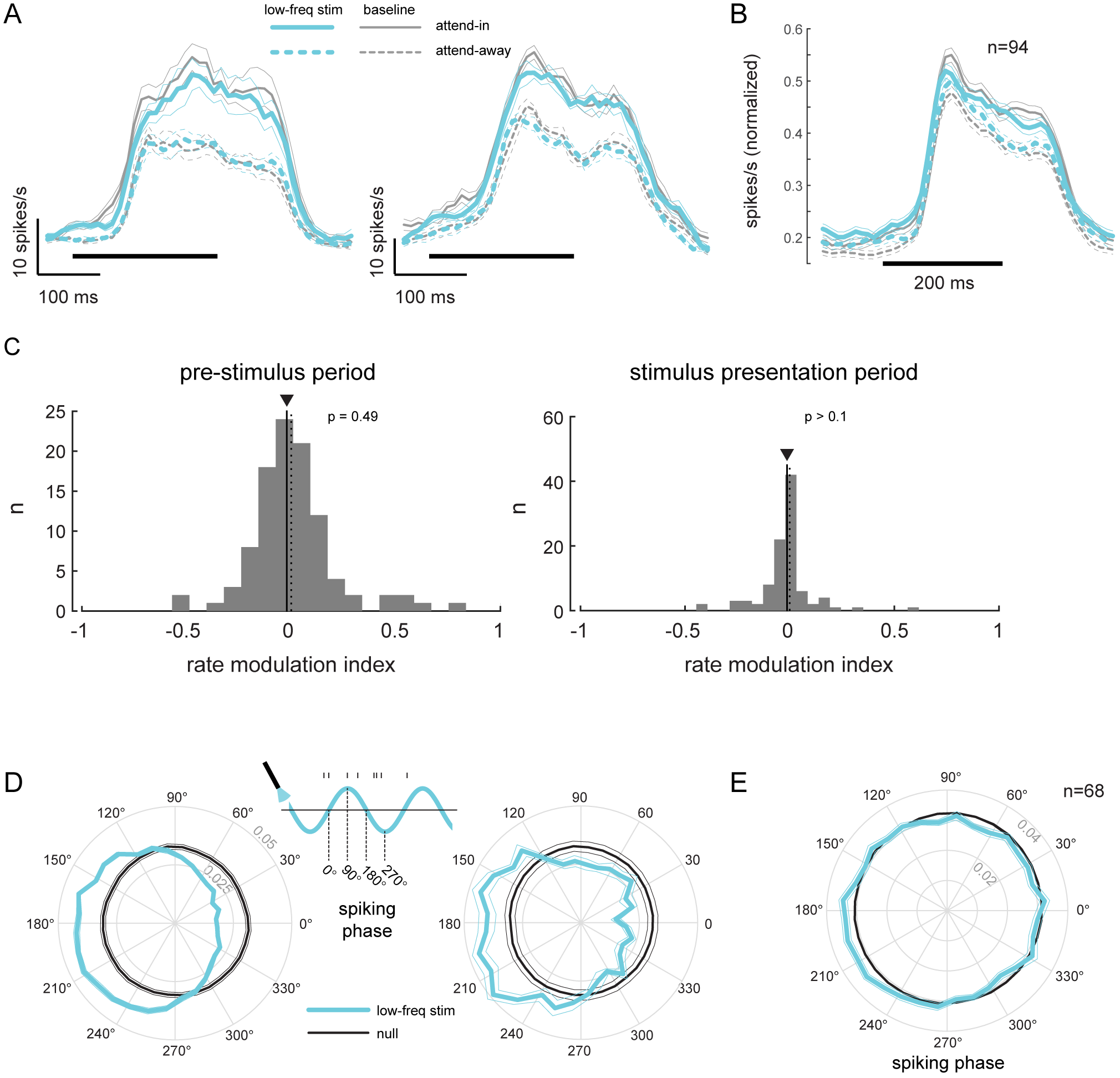
Low-frequency stimulation induces phase-locking without increasing firing rates. (**A**) Peristimulus time histograms (PSTH) of two example units for the different experimental conditions. Both units show a robust firing rate modulation due to attention (solid versus dashed lines) but no rate increase due to low-frequency optical stimulation (blue versus gray lines). Horizontal bars represent stimulus duration. (**B**) Population data showing the same rate increase due to attention, but no significant increase due to optical stimulation (*n* = 94). Same convention as in **A**. (**C**) Distribution of rate modulation indices for the low-frequency stimulation attend-in condition compared to the baseline attend-in condition for a 200ms pre-stimulus period (left panel) and 200ms stimulus presentation period (60-260ms after stimulus onset; right panel). The arrowheads depict the median of the distributions. Neither distribution is significantly different from zero (*p* > 0.1). (**D**) Phase plots for two example units showing the distribution of spiking activity with respect to the phase of the optical stimulation. In gray is the null distribution obtained from a rate-matched Poisson process. Both units show significant deviations from the null distribution (*p* ≪ 0.01 for both, Rayleigh test), indicative of phase locking. (**E**) Population phase-locking plot illustrating the bias in spiking activity to the downswing of optical stimulation (*n* = 68). Same convention as in **D**. The distribution of spiking phase is significantly different from null (*p* < 0.01, Rayleigh test).

Behaviorally, we find that low-frequency stimulation impairs the monkey’s ability to detect fine orientation changes, and does so only in the attend-in condition (Fig 2B, upper panel; Fig 2 - Supp 2B), not in the attend-away condition (Fig 2B, lower panel). To quantify this behavioral deficit, we calculated the threshold and slope of the psychometric functions (see Methods) and assessed changes in these two quantities due to optical stimulation. We find that optical stimulation significantly increases both the threshold (*p* = 0.02, *t*-test) and the slope (*p* = 0.007, *t*-test) of the psychometric function compared to the no stimulation trials in the attend-in condition (2C). In other words, the behavioral curves, on average, are shifted to the right and are steeper, together implying impaired performance.

The impairment due to optical stimulation is location specific: there was no change in performance on trials when the monkey was cued to detect the target at the unstimulated location (attend-away condition, Fig 2D). Importantly, the impairment is also frequency specific. If we stimulate the neurons with 20Hz sinusoidally modulated low-power laser stimulation (‘high-frequency stimulation’ condition), we observe no change in behavior (2E), despite increase in high-frequency spike-count correlations (3B) and phase locking comparable to the low-frequency stimulation condition (Fig 4 - Supp 1).

To confirm whether it is possible to induce coherent activity in a neuronal ensemble due to sub-threshold rhythmic stimulation, we examined the consequences of such stimulation on a conductance-based model of excitatory and inhibitory neurons (Fig 5A; see Methods). We calculated the strength of coherent activity in the network (spike-spike coherence, SSC) both with and without sub-threshold stimulation (Fig 5B,C). We quantified the change in coherence due to stimulation as a modulation index (SSC MI; Fig 5D). We find that it is indeed possible to induce coherent activity in the network at a desired frequency (Fig 5D, Fig 5 - Supp 1D) and that this induction is robust to a wide range of network (Fig 5 - Supp 1C, Fig 5 - Supp 2A) and stimulation parameters (Fig 5 - Supp 2B).

**Fig 5.**
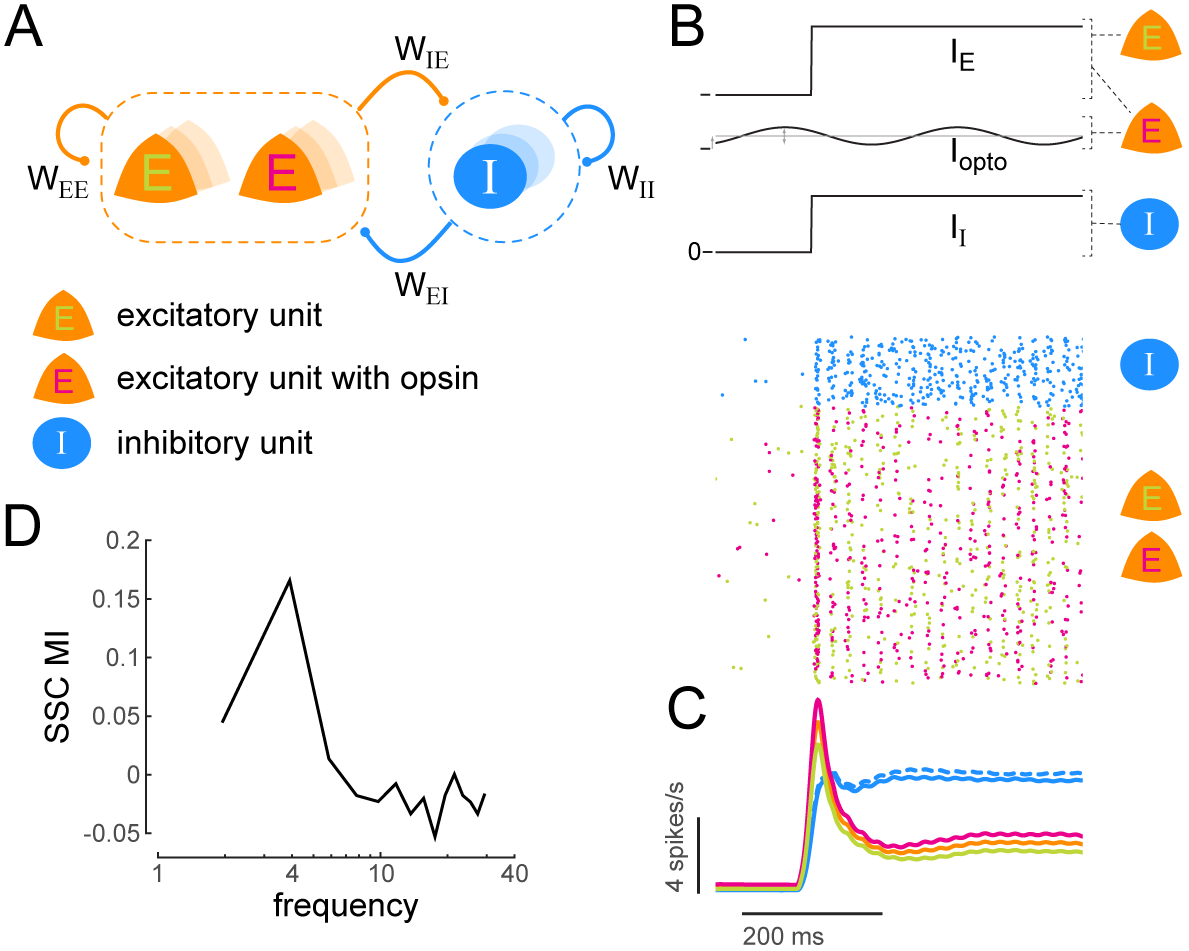
Low-frequency sub-threshold stimulation induces coherent activity in a computational model of E-I neurons. (**A**) Schematic of a local conductance-based E-I network with mutually coupled excitatory (E) and inhibitory (I) units. A fraction (50%) of the E units are sensitive to “optical” stimulation. *W*_*ee*_, self-excitation among E units; *W*_*ii*_, self-inhibition among I units; *W*_*ie*_, excitation provided by E to I; *W*_*ei*_, inhibition provided by I to E. (**B**) Simulation of a network of 800 E and 200 I units (*W*_*ee*_ = 16, *W*_*ii*_ = -1, *W*_*ie*_ = 4, *W*_*ei*_ = -18). The raster plot shows the activity of all units in the model (blue, I; green, E without opsin; magenta, E with opsin) to a step input (*I*_*e*_, *I*_*i*_) and 4Hz sinusoidal optical stimulation (I_opto_). (**C**) Population spiking rate averaged across 1000 simulations of the scenario in **B** with and without optical stimulation. (blue, I; orange, all E; green, E without opsin; magenta, E with opsin. solid lines, with optical stimulation; dashed lines, without optical stimulation) (**D**) Spike‐ spike coherence (SSC) among E units was calculated for the two conditions with and without optical stimulation and the change in SSC across the two conditions was calculated as a modulation index (SSC MI). SSC MI exhibits a peak at 4Hz due to optical stimulation.

The location specificity of the impairment also suggests that the impairment is not due to a phosphene effect (Jazayeri et al., 2012). If attention were drawn away from the unstimulated location by a phosphene we would expect impaired performance in the attend-away condition. We also verified that the impairment was not due to a thermal effect by stimulating a location in the chamber a few millimeters from the opsin site (Fig 2 - Supp 3A) and not due to visual distractions cause by the laser light by stimulating outside the brain (Fig 2 - Supp 3B). In both cases, we did not observe any changes in behavior.

Our results establish the first causal link between correlated variability and perception. The optogenetic stimulation protocol in our study, using sinusoidal modulation of laser irradiance, induces the kind of correlations in a local population of the cortex that might not be physiologically realistic. It nevertheless establishes the causal relevance of low-frequency correlated variability in perception and supports the hypothesis that attention-dependent reductions in correlated variability enhance perception. Recently, studies have theorized that only certain correlations - those that are indistinguishable from stimulus-induced correlations - are information limiting (Moreno-Bote et al., 2014). We speculate that the correlations induced in our study included such information-limiting correlations, resulting in the observed impairment. The timescale of these low-frequency correlations is consistent with inter-saccadic intervals (200-300ms), which may be a relevant timeframe for gathering visual information (Yarbus et al., 1967). Decreases in correlated variability at this timescale could therefore be critical for perception. Our study paves the way for investigating the laminar and cell-class specific components of the cortical circuit that determine this critical component of perception.

## ACKNOWLEDGEMENTS

This research was supported by NIH R01 EY021827 to J.H.R. and A.S.N., the Gatsby Charitable Foundation to J.H.R., fellowship from the NIH T32 EY020503 training grant and the NARSAD Young Investigator Grant to A.S.N., The Salk Institute Excellerators Fellowship Program and the NARSAD Young Investigator Grant to J.J.N. and by a NEI core grant for vision research P30 EY019005 to the Salk Institute. We would like to thank Ed Callaway and Euiseok Kim for help with optogenetic reagents, Rob Teeuwen for assistance with animal training and Catherine Williams and Mat LeBlanc for excellent animal care.

## AUTHOR CONTRIBUTIONS

A.S.N., J.J.N. & J.H.R. designed the experiments. A.S.N. collected and analyzed the data. A.S.N. developed the computational model and ran the simulations. A.S.N., J.H.R. & J.J.N. wrote the manuscript.

## METHODS

### Surgical Procedures

Surgical procedures have been described in detail previously (Nandy et al., 2017; Nassi et al., 2015; Ruiz et al., 2013). In brief, an MRI compatible low-profile titanium chamber was placed over the pre-lunate gyrus, on the basis of preoperative MRI imaging in two rhesus macaques (right hemisphere in Monkey A, left hemisphere in Monkey C). The native dura mater was then removed and a silicone based optically clear artificial dura (AD) was inserted, resulting in an optical window over dorsal V4 (Fig 1A,B). All procedures were approved by the Institutional Animal Care and Use Committee and conformed to NIH guidelines.

### Viral injections

Viral injection procedures have been described in detail previously (Nassi et al., 2015). In brief, we injected a VSVg-pseudotyped lentivirus carrying the C1V1-EYFP gene behind the 1.3kb CaMKIIα promoter (lenti-CaMKIIa-C1V1_E162T_-ts-EYFP; titer = 3x10^10^ TU/ml) at 2 cortical sites in monkey A and 1 cortical site in monkey C while they were anesthetized and secured in a stereotactic frame. The viral constructs were chosen to preferentially drive expression of the depolarizing opsin CiVi in excitatory neurons local to the injection site (Han et al., 2009). We injected approximately 0.5μl of virus at each depth in 200μm increments across the full 2mm thickness of cortex. All injections were targeted to para-foveal regions of V4 with eccentricities between 5 and 8 degrees of visual angle. Expression of the fluorescently tagged opsin was confirmed using epifluorescence goggles (BLS Ltd., Budapest, Hungary) after about 4-6 weeks of viral injection (Fig 1C).

### Opto-Electrophysiology

At the beginning of each recording session, a plastic insert, with an opening for targeting electrodes and for optical stimulation, was lowered into the chamber and secured. This served to stabilize the site against cardiac and respiratory pulsations. The opening was centered at the site of viral injection. A single tungsten microelectrode (FHC Inc.) was mounted on an adjustable X-Y stage attached to the recording chamber and advanced into the injection site using a micromanipulator (Narishige Inc.) until a spike (single neuron or multi-unit) could be reliably isolated from background voltage fluctuations. Site targeting was done under microscopic guidance (Zeiss Inc.) using the microvasculature as reference. A single optical fiber (600 μm multimode fiber, 0.37NA, Thorlabs Inc.) was mounted on the same X-Y stage and positioned over the injection site perpendicular to the calvarium. The microelectrode was positioned at an angle of 20 degrees with respect to the optical fiber (see schematic in Fig 1A).

We used a 532nm diode-pumped solid-state (DPSS) laser (OEM Laser Systems Inc.) as the light source for optical stimulation. The laser was placed on an optical breadboard in-line with a Uniblitz mechanical shutter (Vincent Associates), electrooptical modulator (‘EOM’, ConOptics Inc.) and an optical fiber collimator/coupler (Thorlabs, Inc.) attached to the optical fiber. A beam-splitter between the EOM and collimator directed approximately 1% of the light toward a high-speed photo-detector (Thorlabs, Inc.). The EOM allowed us to control the intensity of laser light entering the fiber and was controlled using custom-written Labview software and a National Instruments digital acquisition board. Before each experiment we calibrated the output of the high-speed photodetector to the full range of intensities (irradiance units) measured at the fiber tip using an integrating sphere photodiode power sensor and a digital power meter (Thorlabs, Inc.). This enabled real-time, calibrated irradiance measurements during all experiments.

Neuronal signals were recorded extracellularly, filtered, and stored using the Multichannel Acquisition Processor system (Plexon Inc.). Neuronal signals were classified as either multi-unit clusters or isolated single units using Plexon Offline Sorter software. Single units were identified based on two criteria: (a) if they formed an identifiable cluster, separate from noise and other units, when projected into the principal components of waveforms recorded on that electrode and (b) if the inter-spike interval (ISI) distribution had a well defined refractory period.

Data was collected over 42 sessions (24 sessions in Monkey A, 18 in Monkey C), yielding a total of 94 units. Frequently, multiple units could be identified while recording from the single tungsten electrodes. Data was collected over an additional 3 sessions for control analyses (Fig 2 - Supp 3).

### Task and Stimuli

Stimuli were presented on a computer monitor placed 57 cm from the eye. Eye position was continuously monitored with an infrared eye tracking system (ISCAN ETL-200). Trials were aborted if eye position deviated more that i° (degree of visual angle, ‘dva’) from fixation. Experimental control was handled by NIMH Cortex software (http://www.cortex.salk.edu).

#### Receptive Field (RF) Mapping

At the beginning of each recording session, neuronal RFs were mapped using subspace reverse correlation (Ringach et al., 1997) in which Gabor (eight orientations, 80% luminance contrast, spatial frequency i.2 cycles/degree, Gaussian half-width 2°) or ring stimuli (80% luminance contrast) appeared at 60 Hz while monkeys maintained fixation. Each stimulus appeared at a random location selected from an iixii grid with i° spacing in the appropriate visual quadrant. All RFs were in the lower visual quadrant (lower-left in Monkey A, lower-right in Monkey C) and with eccentricities between 5 and 8 dva.

#### Irradiance response curves

After estimating the RF of a single-unit or multi-unit cluster, we assessed its sensitivity to optical stimulation. While the monkey maintained fixation, we measured the neuronal response to visual (achromatic Gabor stimulus, spatial frequency i.2 cycles/degree, 20% luminance contrast) and optical stimulation. The visual stimulus was flashed at the RF for 200ms with a simultaneous step laser pulse chosen from one of several irradiance values (typically 0, 10, 30, 50 and 70 mW/mm^2^) (Fig 2 - Supp 4).

#### Attention task

In the main experiment, monkeys had to perform an attention-demanding orientation change-detection task (2A). While the monkey maintained fixation, two achromatic Gabor stimuli (spatial frequency 1.2 cycles/degree, 6 contrasts randomly chosen from an uniform distribution of luminance contrasts, *c* = [10, 18, 26, 34, 42, 50%]) were flashed on for 200ms and off for a variable period chosen from a uniform distribution between 200-400ms. One of the Gabors was flashed in the center of the RFs, the other at a location of equal eccentricity across the vertical meridian. At the beginning of a block of trials, the monkey was spatially cued (‘instruction trials’) to covertly attend to one of these two spatial locations. During these instruction trials, the stimuli were only flashed at the spatially cued location. At an unpredictable time (minimum 1s, maximum 5s, mean 3s), one of the two stimuli changed in orientation. The monkey was rewarded for making a saccade to the location of orientation change. However, the monkey was rewarded for only those saccades where the saccade onset time was within a window of 100-400ms after the onset of the orientation change. The orientation change occurred at the cued location with 95% probability and at the uncued location with 5% probability (‘foil trials’). We controlled task difficulty by varying the degree of orientation change (Δ_orl_), which was randomly chosen from one of 8 orientations in the range 1-15°. The orientation change in the foil trials was fixed at 4°. These foil trials allowed us to assess the extent to which the monkey was using the spatial cue, with the expectation that there would be an impairment in performance and slower reaction times (Fig 2 - Supp 2A) compared to the case in which the change occurred at the cued location. If no change occurred before 5s, the monkey was rewarded for maintaining fixation (‘catch trials’, 13% of trials). We will refer to all stimuli at the baseline orientation as ‘non-targets’ and the stimulus flash with the orientation change as the ‘target’.

On a random subset of trials, neurons at the injection site were stimulated with 45Hz sinusoidally modulated low-power laser stimulation (‘low-frequency stimulation’ condition). The sinusoidal modulation had excursions from a minimum irradiance close to 0 mW/mm^2^ to a maximum irradiance, chosen such that the equivalent root-mean‐ squared intensity elicited a firing rate either 10% above (Fig 2 - Supp 4, left example unit) or 10% below (Fig 2 - Supp 4, right example unit) the firing rate in the zero irradiance condition. The optical stimulation lasted the entire duration of the trial. On a subset of experimental sessions (n=15), neurons at the injection site were also stimulated with 20Hz sinusoidally modulated low-power laser stimulation (‘high‐ frequency stimulation’ condition).

### Data analysis

#### Behavioral Analysis

For each orientation change condition Δ_ori_, we calculated the hit rate as the ratio of the number of trials in which the monkey correctly identified the target with a saccade over the number of trials in which the target was presented. The hit rate as a function of Δ_ori_, yields a behavioral psychometric function (Fig 2B, Fig 2 - Supp 1, Fig 2 - Supp 2). Psychometric functions were fitted with a smooth logistic function (Palamedes MATLAB toolbox). Error bars were obtained by a jackknife procedure (20 jackknives, 5% of trials left out for each jackknife). Performance for the foil trials were calculated similarly as the hit rate for trials in which the orientation change occurred at the un-cued location (Fig 2 - Supp 2A, left panel, square symbol). Performance for the catch trials was calculated as the fraction of trials in which the monkey correctly held fixation for trials in which there was no orientation change (Fig 2 - Supp 2A, left panel, star symbol). Psychometric functions were obtained separately for the baseline (no laser stimulation) and the optical stimulation conditions. We characterized the fitted psychometric functions by two quantities: *threshold*, the stimulus condition at which performance was mid-way between the lower and upper asymptotes; *slope*, the steepness of the psychometric function at threshold. We assessed any behavioral changes due to optical stimulation as changes in threshold and slope with respect to the baseline (no laser stimulation) condition.

#### Peri-stimulus time-histograms

For this and subsequent analyses of neuronal data, we restricted our analyses to non-target flashes from correct trials (hit trials in which the monkey correctly detected a target or correct catch trials). Neuronal responses were binned using a sliding window of width 30ms that was shifted by 10ms increments to obtain the time-varying firing rates, also known as the peri-stimulus time‐ histograms (PSTH), of the recorded units (4A). Population PSTH plots (4B) were obtained after normalizing the responses of each neuron to the peak across the four experimental conditions (2 attention conditions [attend-in, attend-away] x 2 stimulation conditions [no stimulation, laser stimulation]).

#### Spike-phase distributions

We calculated the phase of each spike with respect to the sinusoidal laser stimulation during a 200ms blank period before a non-target stimulus flash. We only considered those inter-stimulus periods where the inter-stimulus interval was greater than 500ms (in other words, the interval between onset of the stimulus and the offset of the previous stimulus was greater than 300ms), so as to minimize artifacts due to stimulus offset. Polar plots in Fig 4C show the distributions of spiking phases. To see if these distributions were significantly different from chance, we calculated a null distribution by generating spike times from a rate-matched Poisson process (gray polar plots in Fig 4C). To obtain reliable estimates for spike-phase distributions, we restricted our analysis to units with a minimum firing rate of 5 spikes/s (n=68; firing rate averaged over the 200ms stimulus-evoked period between 60-260ms after non-target onset).

#### Spike-count correlations (*r*_*SC*_)

We calculated the Pearson correlation of spike counts across trials for every pair of simultaneously recorded units. In order to remove the influence of confounding variables like stimulus strength, spike counts were z‐ scored using the mean and standard deviation for repetitions of each stimulus type. Ordered pairs of z-scored spike counts were collapsed across contrast conditions and the Pearson correlation was calculated from these ordered pairs. This was done separately for the different attention and optical stimulation conditions and also for different sized counting windows (50ms for high-frequency correlations, 200ms for low‐frequency correlations) during the stimulus-evoked period between 60-260ms after nontarget onset (Fig 3). Multiple non-overlapping windows were used for those counting windows that were smaller than the 200ms stimulus evoked period.

### Computational model

A similar model has been described previously (Nandy et al., 2017). We set up a conductance-based model of *N*_E_ excitatory and *N*_I_ inhibitory neurons with 80% connection probability (both within and across the two populations) and with the following synaptic weights (5A):

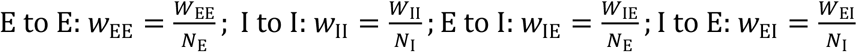

We simulated models of *p*_E_ = 800 excitatory and *p*_I_ = 200 inhibitory spiking units. The spiking units were modeled as Izhikevich neurons (Izhikevich, 2003) with the following dynamics:

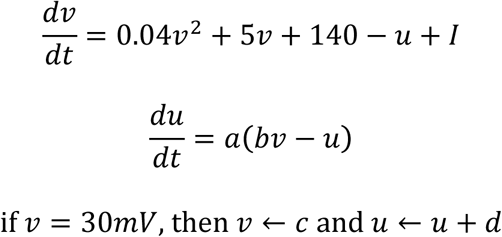

*v* is the membrane potential of the neuron and *u* is a membrane recovery variable. *I* is 2 the current input to the neuron (synaptic and injected DC currents). The parameters *a*, *b*, *c* and *d* determine intrinsic firing patterns and were chosen as follows:

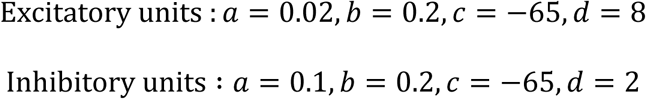

Presynaptic spikes from excitatory units generated fast (AMPA) and slow (NMDA) synaptic currents, while presynaptic spikes from inhibitory units generated fast GABA currents:

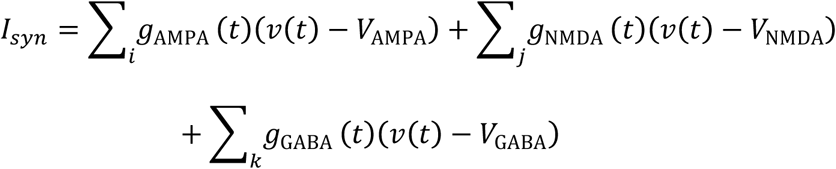

where *V*_AMPA_ = 0, *V* _NMDA_ = 0, *V*_GABA_ = -70 are the respective reversal potentials (mV). 8 The synaptic time courses *g*(*t*) were modeled as a difference of exponentials:

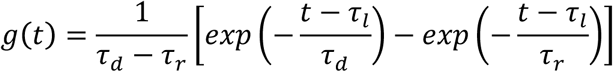

where *tau*_*l*_,*t*_*r*_ and *tau;*_*d*_ are the latency, rise and decay time constants with the following 10 parameter values (ms) (Brunel and Wang, 2003):

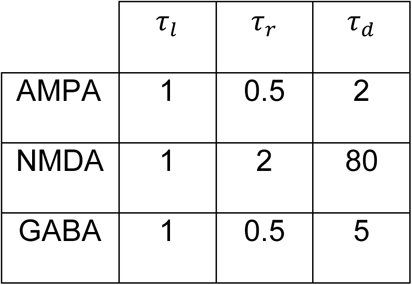

The NMDA to AMPA ratio was chosen as 0.45 (Myme et al., 2003).

The network was stimulated by a DC step current (*I*_E_ = 2.8, *I*_*I*_ = 2.3) of duration 1.5s (5B). Synaptic noise was simulated by drawing from a normal distribution (*I*_syn-noise_∼*N*(*μ* = 0, |σ = 3). To simulate the laser stimulation in the main experiment, we chose a random subset (50%) of excitatory units to which we injected a 4Hz sinusoidally modulated current (*I*_opto_;mean current = 0.5; peak to trough range = 0.7). Such a current by itself did not produce spiking activity in the network.

We computed the spike-spike coherence between all pairs of excitatory units in the model (irrespective of whether the units were subjected to the additional sinusoidally modulated current) using multi-taper methods (Mitra and Pesaran, 1999), over a 400ms window for both simulation conditions: with and without *I*_opto_. Spike trains were tapered with a single Slepian taper, giving an effective smoothing of 2.5Hz for the 400ms window (NW=1, K=1). To control for differences in firing rate between the two conditions, we adopted a rate matching procedure similar to (Mitchell et al., 2009). Induction of coherent activity in the network due to sub-threshold sinusoidal stimulation was calculated as a modulation index of coherence across the two conditions: SSC MI = (SSC_with_-SSC_without_)/(SSC_with_+SSC_without_). In order to obtain a baseline for the coherence expected solely due to trends in firing time-locked to network stimulation, we also computed coherence in which trial identities were randomly shuffled (Fig 5 - Supp 1C-D).

---

**Fig 2 - Supp 1.**
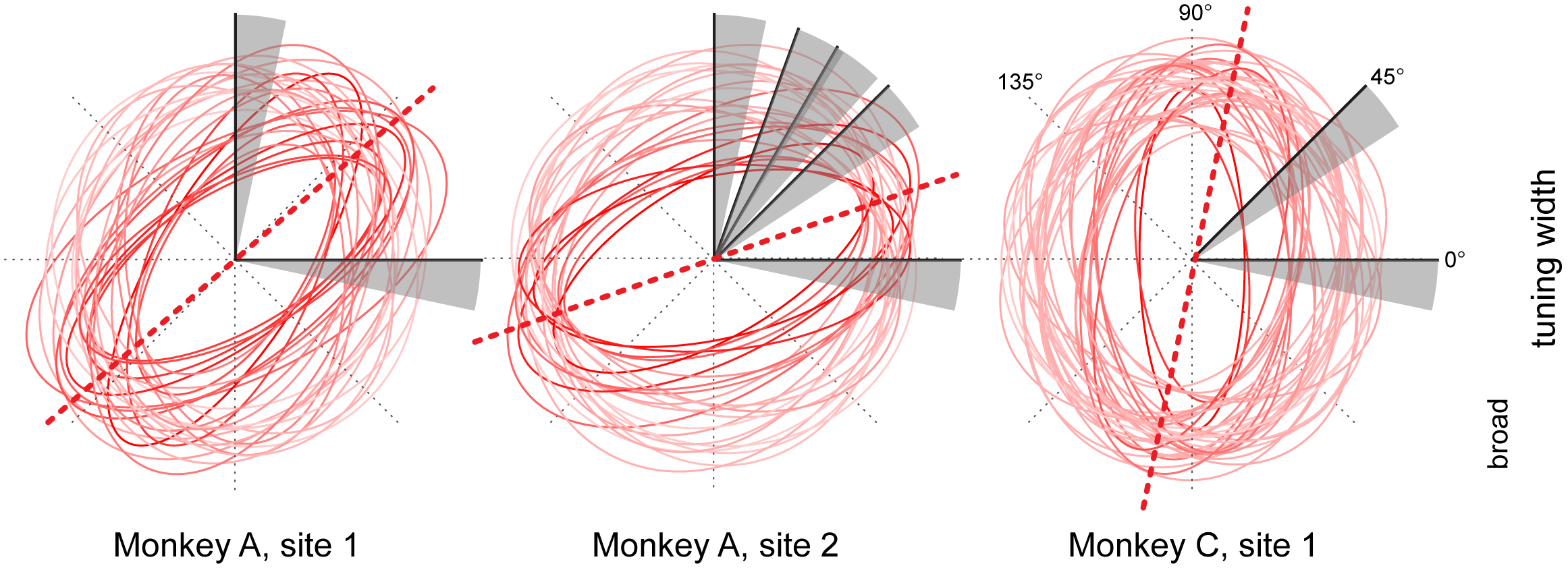
Orientation tuning properties at opsin injection sites. Orientation tuning plots of single units and multi-unit activity at the different cortical injection sites (2 in monkey A, 1 in monkey C) were fitted with ellipses (least-squares fit). The fitted ellipses are overlaid to illustrate the overlap in tuning at each injection site. The ellipses are color-coded such that more saturated colors correspond to units with sharper orientation tuning, as estimated from the aspect ratio of the fitted ellipse. The dotted red lines represent the average peak tuning at each site. The gray sectors represent the range of orientation changes that the monkeys had to detect with respect to baseline orientation (black line within each sector) across different behavioral sessions.

**Fig 2 - Supp 2.**
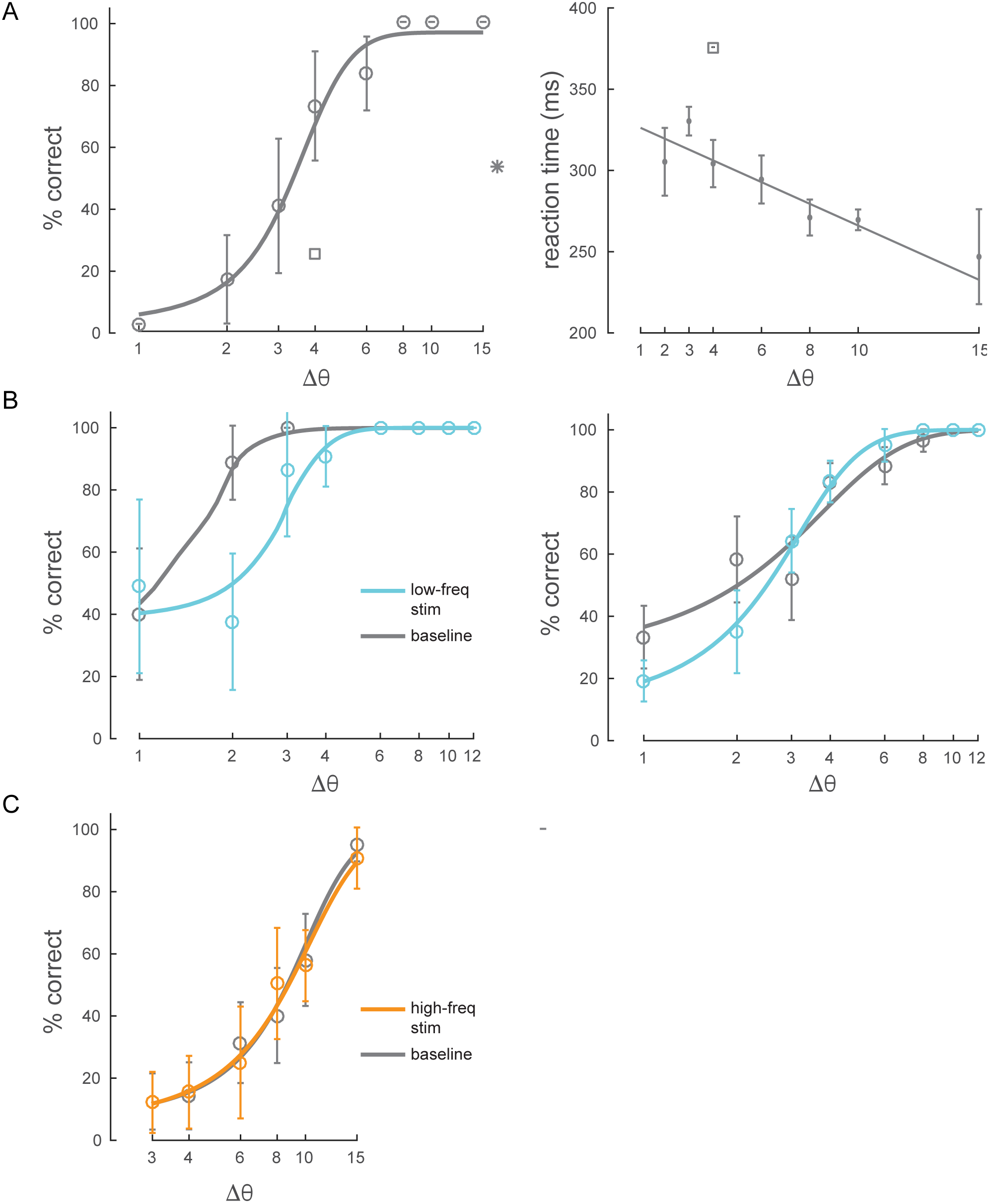
Behavioral performance. (**A**) *Left panel*: example behavioral session showing performance (hit rate) as a function of task difficulty (size of orientation change) for the baseline condition (no optical stimulation). Square symbol: foil-trial performance. Star symbol: catch-trial performance. Error bars are std. dev. obtained by a jackknife procedure and corrected for the number of jackknives (20). The data has been fitted with a smooth logistic function. *Right panel*: reaction time as a function of task difficulty for the same session. The data has been fitted with a linear regression line. Performance is degraded and reaction times are slower for the foil trials, indicating that the animal was indeed deploying attention to the spatially cued location. (**B**) Two example sessions showing a behavioral impairment due to low-frequency (4-5Hz) optical stimulation when the monkey was attending in to the site of optical stimulation. Same format as in Fig 2B, top panel. (**C**) Example behavioral session showing no change in performance due to high-frequency (20Hz) optical stimulation.

**Fig 2 - Supp 3.**
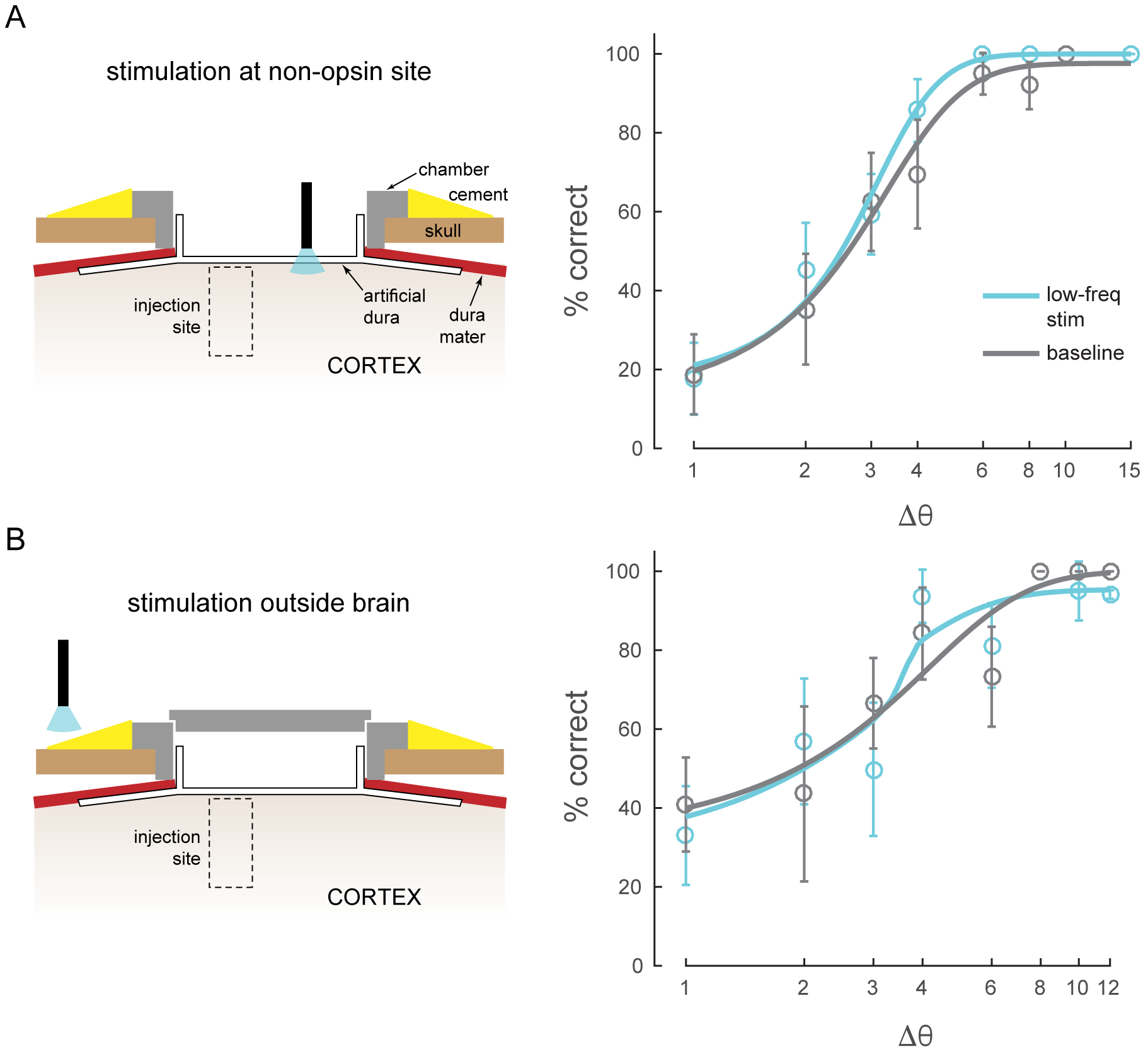
Other control conditions. (**A**) Stimulation at a non-opsin site does not perturb behavior. (**B**) Stimulation with optical fiber outside the brain (with the opto-physiology chamber closed) does not perturb behavior.

**Fig 2 - Supp 4.**
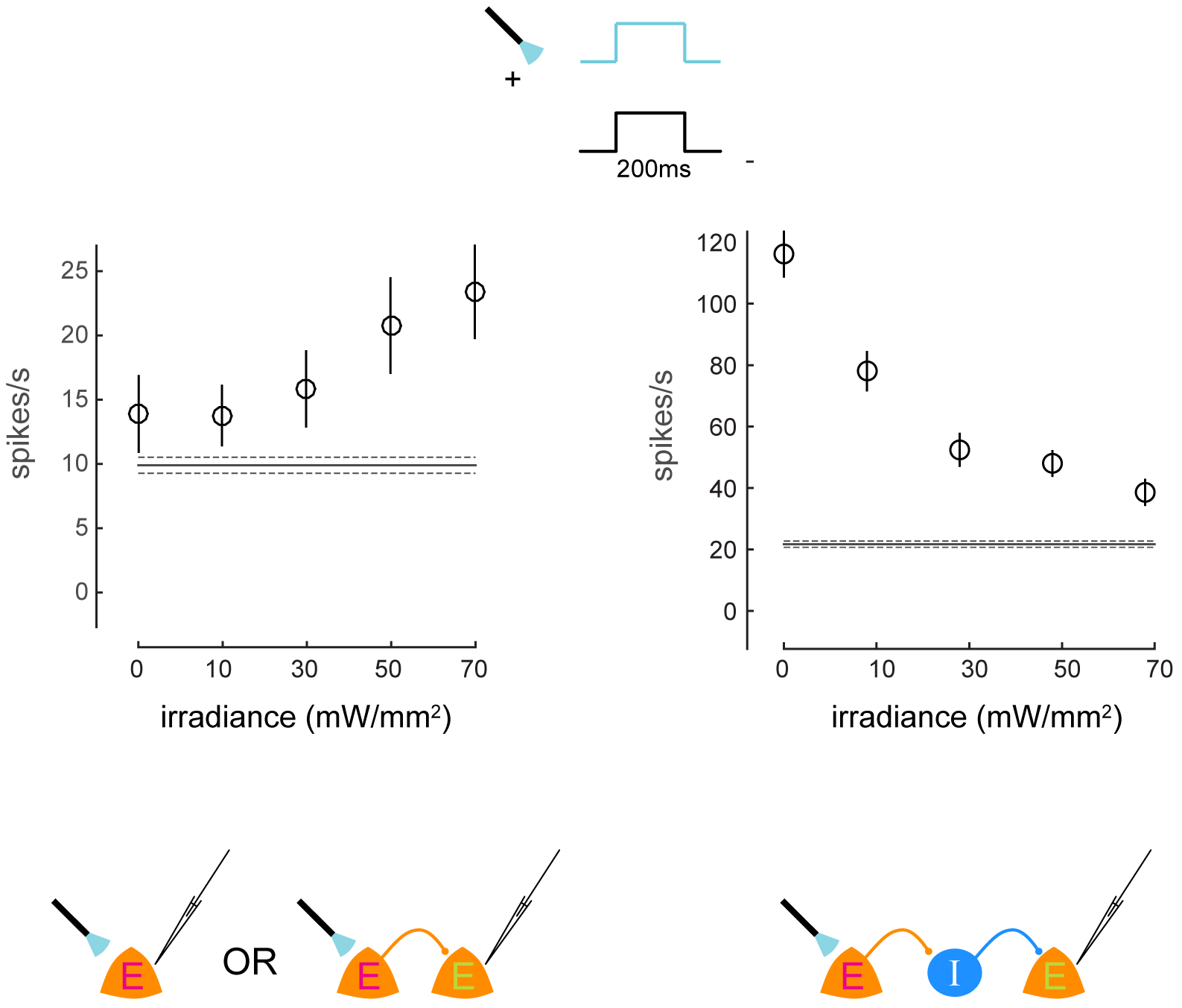
Irradiance response curves. Response of two neurons to presentations of a visual stimulus and simultaneous optical stimulation. The visual stimulus was a 20% contrast Gabor stimulus presented for 200ms. The optical stimulation consisted of a concurrent light pulse of various irradiance values (x-axis). The first unit exhibits increases in firing rate with increasing intensity of optical stimulation. The second unit exhibits decreases in firing rate with increasing intensity of optical stimulation, due to indirect network effects. Schematics at the bottom illustrate possible network scenarios. E=excitatory neuron (magenta E = opsin expressing; green E = non-opsin expressing). I=inhibitory neuron. The horizontal line represents baseline firing-rate. Mean +/- s.e.m. in all plots.

**Fig 4 - Supp 1.**
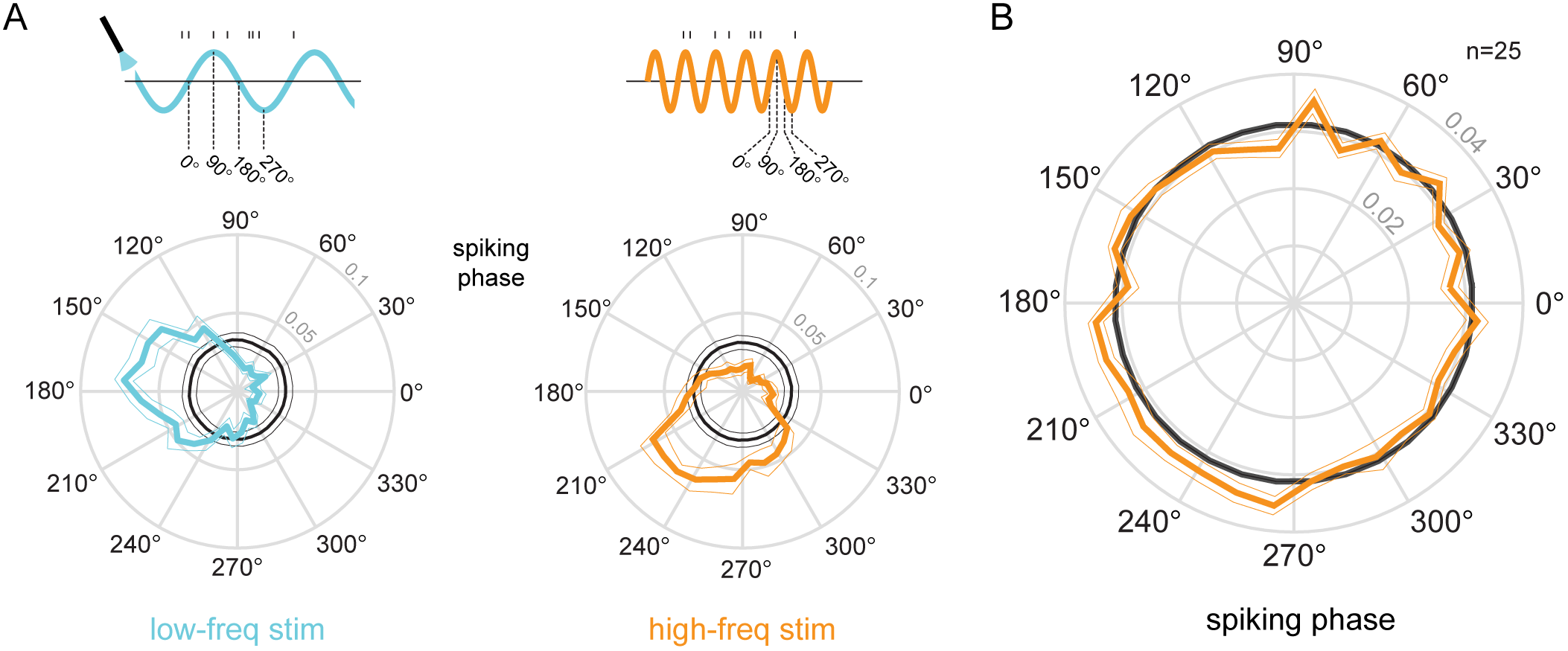
Phase-locking to optical stimulation. (**A**) Phase plots for an example unit showing the distribution of spiking activity with respect to the phase of the low-frequency (5Hz, *left*) and high‐frequency (20Hz, *right*) optical stimulation. In gray is the null distribution obtained from a rate-matched Poisson process. The unit shows significant deviations from the null distribution, indicative of phase locking to both stimulations (*p* ≪ 0.01 for both stimulation conditions, Rayleigh test). (**B**) Population phase-locking plot illustrating the bias in spiking activity to the high-frequency optical stimulation (n = 25). Same convention as in **A**.

**Fig 5 - Supp 1.**
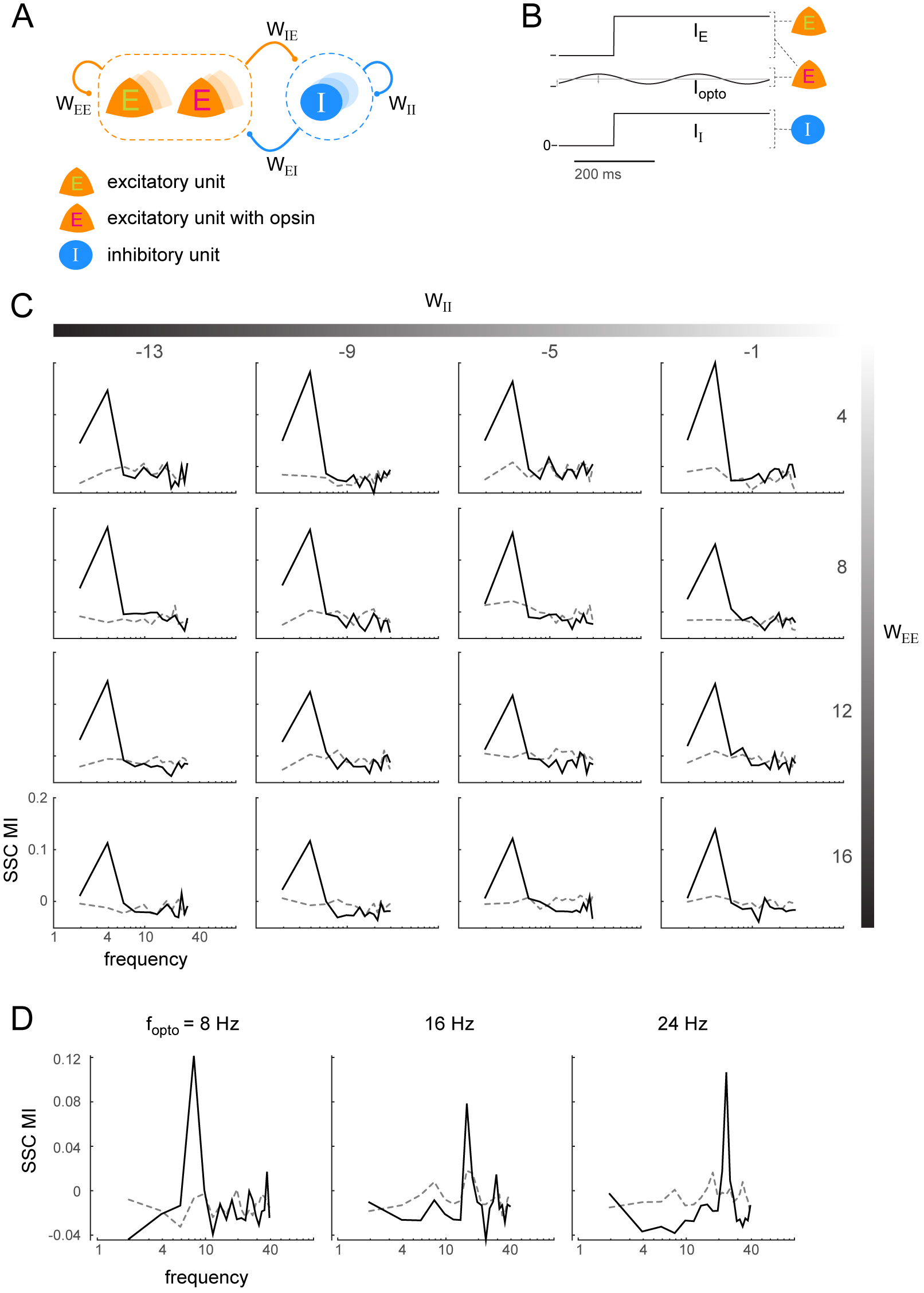
Induction of coherent activity in the E-I model is robust across network and stimulation parameters - I. (**A**)-(**B**) same as Fig 5A-B. (**C**) Spike-spike coherence modulation index (SSC MI; see Fig 5D) as a function of varying the self-coupling parameters *W*_*ie*_ and *W*_*ii*_, keeping the other two parameters fixed (*W*_*ie*_ = 4, *W*_*ei*_ = -18). SSC MI exhibits a peak at the 4Hz optical stimulation frequency irrespective of the self-coupling parameters. The dotted lines represent the null SSC MI obtained by shuffling trial identities. (**D**) The peak SSC MI shifts with increasing optical stimulation frequency (*f*_opto_, network parameters same as in Fig 5), indicating that the peak is not due to any intrinsic resonant activity in the network. Dotted lines same as in **C**.

**Fig 5 - Supp 2.**
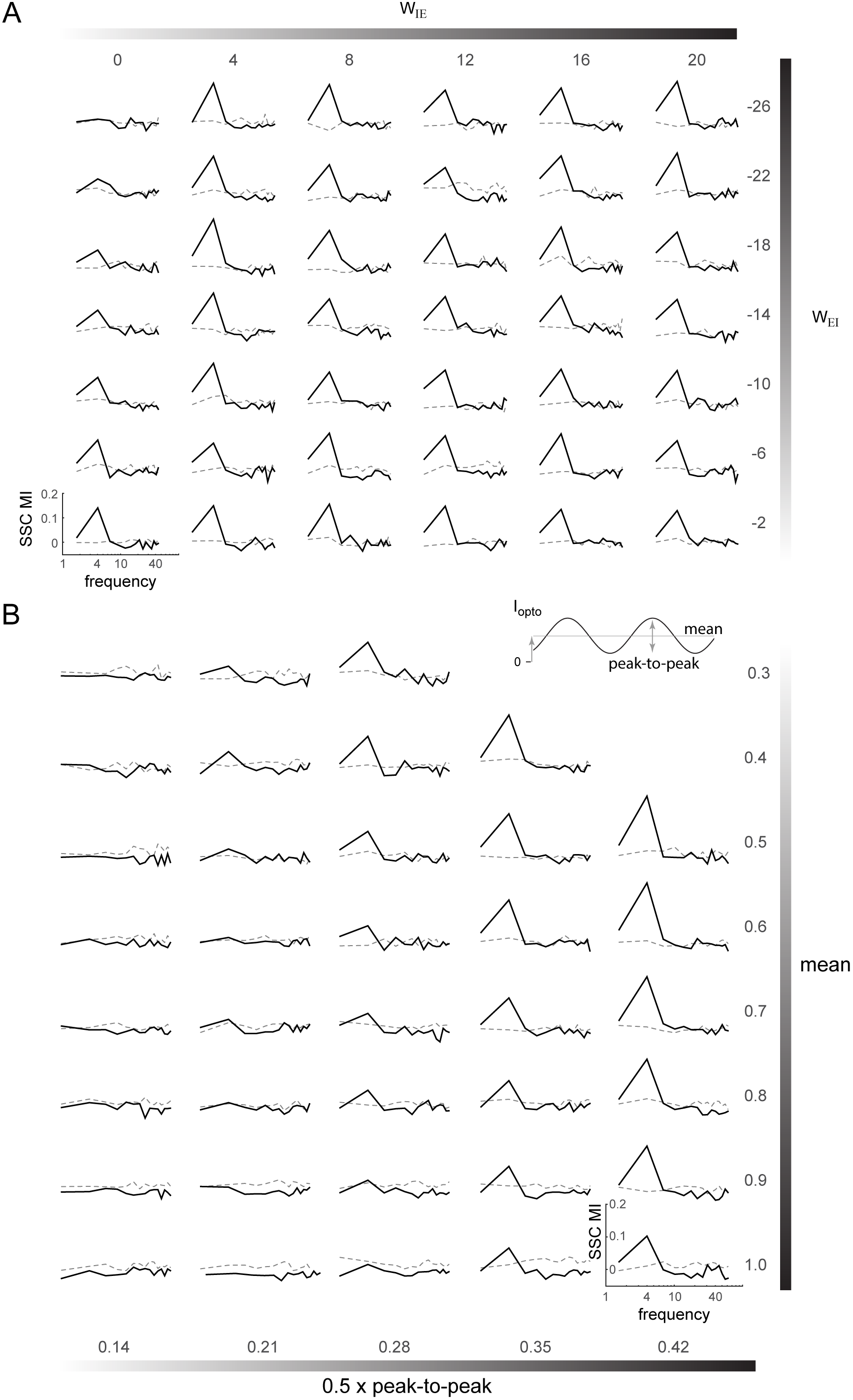
Induction of coherent activity in the E-I model is robust across network and stimulation parameters - II. (**A**) Spike-spike coherence modulation index (SSC MI; see Fig 5D) as a function of varying the cross-coupling parameters *W*_*ie*_ and *W*_*ei*_, keeping the other two parameters fixed (*W*_*ee*_ = 16, *W*_*ii*_ =-1). (**B**) SSC MI as a function of the mean and peak-to-peak variation of the 4Hz sinusoidal optical stimulation (*I*_opto_). Network parameters same as in Fig 5.

